# Parallel circadian-like oscillations in LTP and excitation inhibition balance in mouse CA1 reverse direction after puberty

**DOI:** 10.1101/2025.10.15.682451

**Authors:** Gonzalo Valdivia, Cristian Moreno, Kaiwen He, Darwin Contreras, Trinh Tran, Anthony D. Ramnaugh, William Xu, Altagracia Contreras, Diego C. Fernandez, Daniel Severin, Samer Hattar, Michela Gallagher, Alfredo Kirkwood

## Abstract

Long-term potentiation (LTP), the best-characterized form of Hebbian synaptic plasticity, is well known to be under strong circadian regulation. In mice and rats, both nocturnal species, most studies indicate that LTP in the hippocampal CA1 region is more robust when induced during the dark phase. Our examination of the underlying mechanisms at the CA3 to CA1 synapse provides evidence that the capacity to express LTP does not differ between the light and dark cycles of the 24-hour day. Instead, the magnitude of theta-burst stimulation–induced LTP (TBS-LTP) correlates with daily fluctuations in the ratio of synaptic excitation to inhibition (E/I ratio): both the E/I ratio and TBS-LTP are higher during the dark phase. Consistent with a causal relationship, blockade of inhibition abolishes the light–dark difference in TBS-LTP induction, likewise, pairing-induced LTP, which is less constrained by inhibitory recruitment, does not differ between cycles. Supporting this model, using the APP/PS1 model of AD we observed that neither the E/I ratio nor TBS-LTP change during the light cycle. Finally, we made the intriguing observation that these daily oscillations reverse direction after puberty in WT mice, shifting from being larger in the dark cycle of 2-month-old mice to being larger in the light cycle in 8-month-old mice. This developmental switch may reflect an age-dependent reorganization of circadian control over hippocampal plasticity.

## INTRODUCTION

Long-term potentiation and depression (LTP and LTD, respectively) are extensively studied as models of Hebbian synaptic plasticity that drive network remodeling necessary for learning and memory. Because learning contingencies are highly variable, research on Hebbian plasticity examines not only its mechanisms but also how it is regulated. A current core consensus for LTP/LTD mechanisms is that they are initiated by postsynaptic Ca²⁺ influx through NMDA receptors, followed by the activation of protein kinases, and culminating in the trapping of AMPA receptors at the synapse (for reviews see (Bayer and Giese, 2025; Caya-Bissonnette and Beique, 2024; Lomo, 2025; Nowacka et al., 2024)). Concerning its regulation, it is well established that Hebbian plasticity undergoes postnatal developmental control, characterized by distinct changes in NMDAR subunit composition (Smith et al., 2009; Yashiro and Philpot, 2008)). Hebbian plasticity is also regulated by neuromodulators associated with behavioral states via phosphorylation of NMDAR and AMPA receptors ((Huang et al., 2012; Lutzu and Castillo, 2021; Mihalas et al., 2021; Seol et al., 2007)). Another variable known to strongly influence the magnitude of synaptic plasticity is the time of day, yet the underlying mechanisms remain unclear.

Over four decades ago, Harris and Teyler (Harris and Teyler, 1983) reported that the magnitude of LTP evoked in rat hippocampal slices from CA1 and dentate gyrus regions, differs between the light and dark phases of the day. Similar circadian-like regulation of hippocampal LTP has been documented *in vivo* (Bowden et al., 2012), as well as in mice (Besing et al., 2017; Chaudhury et al., 2005; Dana and Martinez, 1984; McCauley et al., 2020) and hamsters (Raghavan et al., 1999). In addition, cortical LTP, like in CA1, is greater during the dark phase (Alfonsa et al., 2025). In addition, compelling indirect evidence from *in vivo* studies suggests that cortical synaptic potentiation occurs largely during this phase (Hanlon et al., 2011; Miller et al., 2022; Vyazovskiy et al., 2008). The mechanisms that regulate LTP during the day are not fully clear, with studies implicating changes in the NMDAR receptors (Barone et al., 2023) or shifts in the reversal potential of GABA responses (Alfonsa et al., 2025). Our re-examination of the issue at the CA1 to CA3 synapse identifies the daily oscillation of the excitatory/inhibitory balance, measured as a ratio (E/I ratio), as a key daily regulator of LTP induction. Importantly, our results revealed a post-pubertal reversal of this regulation: whereas the E/I ratio and LTP are higher during the dark phase in young adults, they become more prominent during the light phase at later ages. We propose that this age-dependent reversal may underlie some of the discrepancies reported in the field.

## MATERIALS AND METHODS

### Animals

Young (1 and 2-month-old) and adult (8-month-old) C57BL/6 mice as well as 6-month old APPswe;PS1ΔE9 Tg along their wild-type (WT) littermate mice (129/C57BL/6 mixed background) of both sexes were used. APPswe;PS1ΔE9 Tg mice have accelerated amyloid pathologies and have a substantial number of plaque deposits by 6 months of age (Jankowsky et al., 2004; Savonenko et al., 2005). All procedures performed with these animals were approved by the Institutional Animal Care and Use Committees of Johns Hopkins University. Animals were kept at a controlled temperature (22 ± 1 ° C), in light/dark cycles (12:12) with water and food *ad libitum*. Prior to the experiments, mice were entrained for at least two weeks to a 12:12 light:dark cycle in custom entrainment chambers, with the timing of lights-on adjusted according to the chosen circadian time of study (*Zeitgeber* time, ZT = 0, 6, 12 or 18 hours). Mice scheduled for ZT0 and ZT12 were sacrificed 15 minutes before lights on and lights off, respectively. Thus, ZT0 and ZT12 correspond to the end of the dark and light phase, respectively

### Synaptic responses

Hippocampal slices (300 and 400 μm) were prepared as described previously (Ardiles et al., 2012; Boric et al., 2008; Megill et al., 2015). For mice older -weeks , before isolating the brain, the subjects were transcardially perfused, under isoflurane anesthesia with ice-cold dissection buffer containing: 212.7 mM sucrose, 2.6 mM KCl, 1.23 mM NaH2PO4, 26mM NaHCO3, 10 mM dextrose, 3mM MgCl2, and 1mM CaCl2 bubbled with a mixture of 5% CO2 and 95% O2 and recovered at room temperature in

Artificial Spinal Cerebral Fluid (ACSF) that replaced the sucrose by 124 mM NaCl. For Synaptic responses were evoked with 0.2 ms pulses (10 – 80 μA) delivered through concentric bipolar theta glass micropipettes filled with ACSF placed in the stratum radiatum. The baseline rate was 0.033 Hz in all cases. Responses were monitored either extracellularly as field potentials in the stratum radiatum or intracellularly with whole cell-recordings in the soma.

### Field potential recordings

Synaptic responses were evoked with 0.2 ms pulses (10 – 80 μA) delivered through bipolar theta glass micropipettes filled with ACSF, recorded extracellularly in CA1, and quantified as the initial slope of the field potential. Baseline responses were recorded at 0.033 Hz using a stimulation intensity that evoked a half-maximal response, defined as the maximal response without a population spike (pop-spike). Slices were discarded when the pop-spike appeared in the initial rising phase (an indication of hyperexcitability), when paired-pulse facilitation at a 50 ms interval was less than ∼10% (i.e., response 2/response 1 <1.1), or when the baseline was not stable (∼5% drift). LTP was induced using a theta burst stimulation[TBS; four trains, each consisting of ten 100 Hz bursts (four pulses) given at 5 Hz, repeated at 10 s intervals (4×TBS)]. LTP was quantified as the % change the initial slope data and it was expressed as means ± SEM. In experiments including the GABAa antagonist Gabazine (5μM), the CA3 was removed to minimize reverberating activity.

### Whole-Cell Recordings

Visualized whole-cell voltage clamp recordings were made from pyramidal neurons in L2/3 (35% depth from the pia) of V1. Glass pipette recording electrodes (3–6 MΩ) were filled 8 mM KCl, 125 mM cesium gluconate, 10 mM HEPES, 1 mM EGTA, 4 mM Mg-ATP, 0.5 mM Na-GTP, and 5 mM QX-314, and adjusted to pH 7.2–7.3, 280–295 mOsm. Cells with an input resistance ≥ 150 MΩ and access resistance ≤ 25 MΩ were recorded. For all whole cell recordings, cells were discarded if these values changed more than 25% during the experiment. Data were filtered at 2 kHz and digitized at 10 kHz using Igor Pro (WaveMetrics, Portland, OR).

To determine the excitation/inhibition ratio, current responses were recorded in the presence of 100 μM DL-APV. Reversal potentials for excitatory and inhibitory currents of +10 mV and −55 mV (without junction potential compensation) were used (Bridi et al., 2020). A series of stimulations over a range of intensities was delivered, and the responses producing a stable E/I ratio were used (Bridi et al., 2020). Pairing-induced LTP was performed in voltage clamp mode by delivering 150 stimuli at 1 Hz while holding the cell at 0 mV. Changes in synaptic strength were quantified as changes in the current response amplitude recorded at −70 mV.

NMDA/AMPA ratios were calculated from excitatory currents evoked electrical stimulation. Two times the stimulus intensity that evoked the minimal AMPA response was used. An internal solution containing (in millimolar) 102 cesium gluconate, 5 TEA chloride, 3.7 NaCl, 20 Hepes, 0.3 Na-GTP, 4 Mg-ATP, 0.2 EGTA, 10 BAPTA, 5 QX-314 (pH 7.2, ∼300 mOsm) under voltage–clamp (Vh = −70 mV for AMPA and Vh = +40 mV for NMDA). To isolate glutamate evoked currents, ACSF in the recording chamber contained 2.5 μM gabazine.

### General Activity Measurements

General activity was measured using infrared motion detectors from Mini Mitter (Respironics). Mice were housed individually, and the motion detector was mounted on top of the cage to continuously monitor locomotor activity. Data were collected in 10-minute bins using VitalView software (Respironics). Circadian period length was determined by fitting a regression line to the daily onsets of activity over a 7-day period using ClockLab (Actimetrics).

Initially, animals were kept in a standard 12:12 light:dark (LD) cycle, and both total activity and activity during the inactive (light) phase were measured. A 6-hour phase advance was then introduced to assess the phase-shift response, calculated as the number of days required for animals to re-align their activity onset with the new lights-off time. The cycle was subsequently returned to the original phase, this time with a 6-hour delay, to evaluate the masking response—defined as the acute suppressive effect of light on locomotor activity during hours previously corresponding to darkness. Finally, the animals were exposed to constant darkness (DD) to determine their intrinsic circadian period, or free-running rhythm.

## RESULTS

### Daily oscillations of LTP and LTD

The daily regulation of LTP is well established, yet much less is known about LTD, its complementary Hebbian counterpart. Evidence is limited to a single study in anesthetized rats, which reported no LTD during the dark phase (Yang et al., 2012). Therefore, we examined both LTP and LTD in the CA3→CA1 path of slices harvested at different time points during the light/dark cycle (ZT times, see methods). First, we compared LTP and LTD of field responses induced with patterned extracellular stimulation protocols known to be highly effective: theta burst stimulation (TBS) for LTP, low frequency stimulation (LFS) for LTD. The magnitude of TBS-induced LTP (averaged between 50 to 60 min post conditioning) was larger in slices harvested at the end of the dark cycle, ZT0 (see methods), than in slices collected at the end of the light cycle, ZT12, (t-test; p=0.0001. Figure 1A). In the case of LFS-induced LTD, there were no significant differences at these two ZTs (t-test: p=0.7810. Figure 1B). We considered the possibility that the absence of time-of-day differences in LTD was due to inadequate sampled times (ZT0 and ZT12). For example, TBS-LTP and LFS-LTD might both oscillate during the day, but out of phase. Therefore, we collected additional LTP and LTD data at ZT6 and ZT18 times. TBS-LTP magnitude clearly oscillated during the day (ANOVA: F[3,93]=7.705, p=0.001) with ZT12 being smaller than ZT0 and ZT6 (Sidak’s test). In contrast, there was no statistical difference for LFS-LTD evoked at these ZT moments (ANOVA: F[3,78]=0.1196, p=0.9483). The reduced magnitude of TBS-LTP at ZT12 prompted us to evaluate in ZT0 and ZT12 slices the synaptic changes induced with a wide range of stimulation frequencies (1-10 Hz, 900 pulses; 50-100 Hz 300 pulses). The frequency-dependence curves for plasticity induction revealed clear differences between both ZT times (2-way ANOVA. Freq: F[1,96]=14.41, p=0,003; ZTtime: F[3,94]28.97, p<0.0001; Interaction: F[3,94]=4.0054, p=0.0054), particularly at the LTP-inducing range (100Hz, Sidak’s multiple test), with an overall modest potentiation at ZT0. Altogether, the results support the notion that the daily regulation of plasticity affects more LTP than LTD. In addition, our results indicate that the intrinsic capacity to support LTD is present during the dark cycle, and its absence in anesthetized rats may reflect external influences such as neuromodulatory tone (Yang et al., 2012).

**Figure 1.**
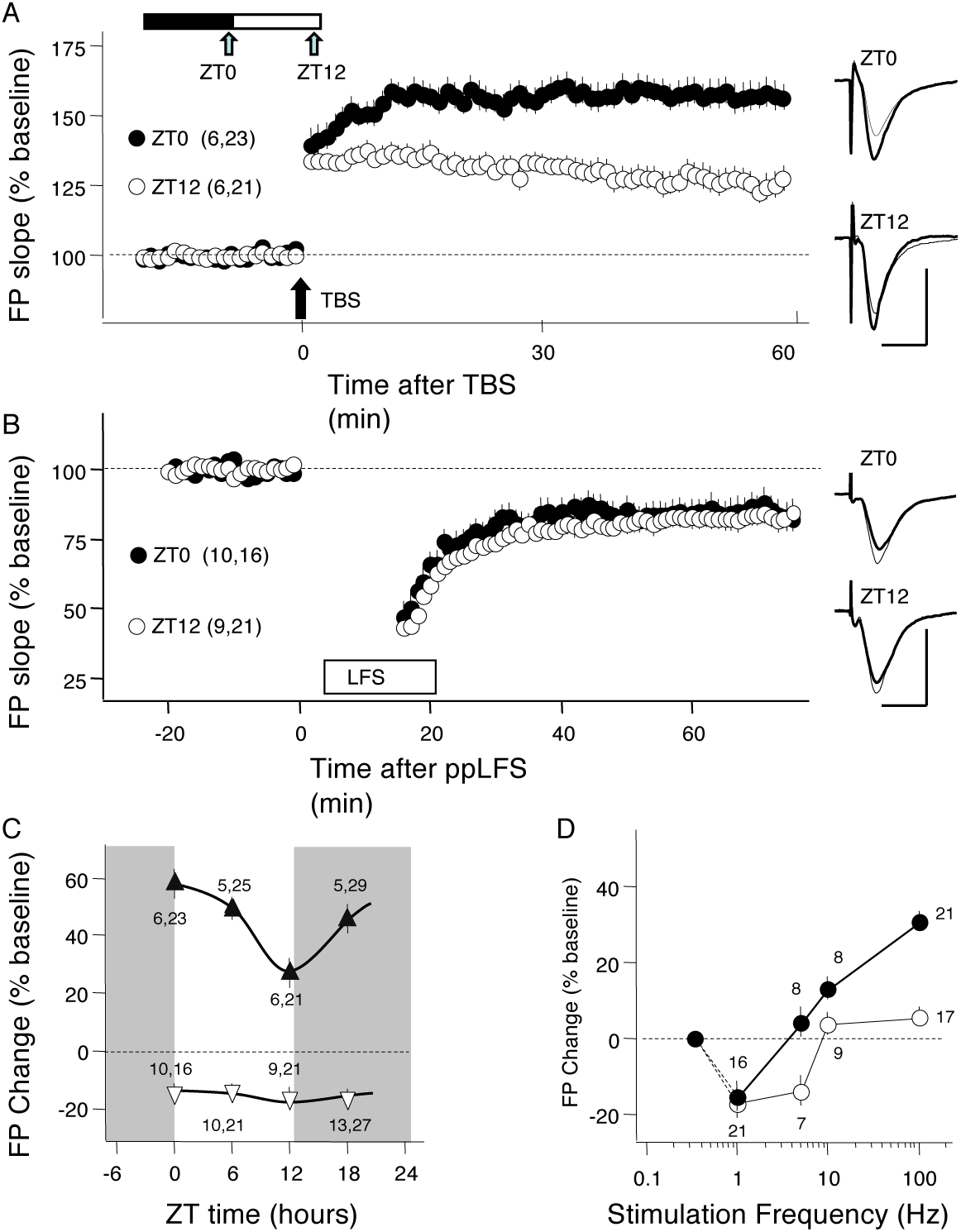
Daily oscillations in the induction of LTP, but not LTD, in CA3→CA1 synapses. A-B) Time course of the changes in FP slope induced by either TBS (A), or LFS (B) (see methods for details) in slices harvested near ZT0 (black circles) or ZT12 (open circles). In each case, the example traces on the right are averages of 4 consecutive responses recorded before (thin lines) and 60 min after conditioning (thick) lines. In parenthesis is the number of mice and slices. C) Average change in FP slope induced by either TBS (black triangles) or LFS (open triangles) in slices harvested at the indicated ZT times. The connecting lines were drawn for visual purposes. D) Plotted is the frequency-dependence function of the average changes in FP slope induced in slices harvested at ZT0 (black circles) and ZT12 (open circles). Indicated in each point is the number of slices included. The grey square represents the absence of change at baseline frequency. Data points in A-D represent average±s.e.m. In A-C, the number of mice and slices is indicated in parenthesis.

### Excitation/inhibition ratio regulates TBS-LTP

Subsequently we examined plausible biophysical mechanisms that could constrain TBS-LTP at ZT12. As candidate mechanism we considered daily reductions in presynaptic release, in synaptic NMDAR function and an enhanced recruitment of synaptic inhibition. We evaluated presynaptic function by determining paired-pulse facilitation (PPF) of the field potentials (see methods). The paired-response ratio, commonly used to detect changes in release probability, was no different between ZT0 and ZT12 at all inter-stimulus intervals tested (Figure 2A. 2-way ANOVA: Interval: F (6, 259) = 59.62, p=0,0001; ZT time: F (1, 259) = 1.501, p<0.22161; Interaction: F (6, 259) = 0.9096). To evaluate changes in NMDAR function we computed the ratio of the AMPAR-mediated and NMDAR-mediated components of the evoked EPSC (see methods). Again, the AMPAR/NMDAR response ratio was no different between ZT0 and ZT12 slices (Figure 2B, t-test: p=0.6095).

**Figure 2.**
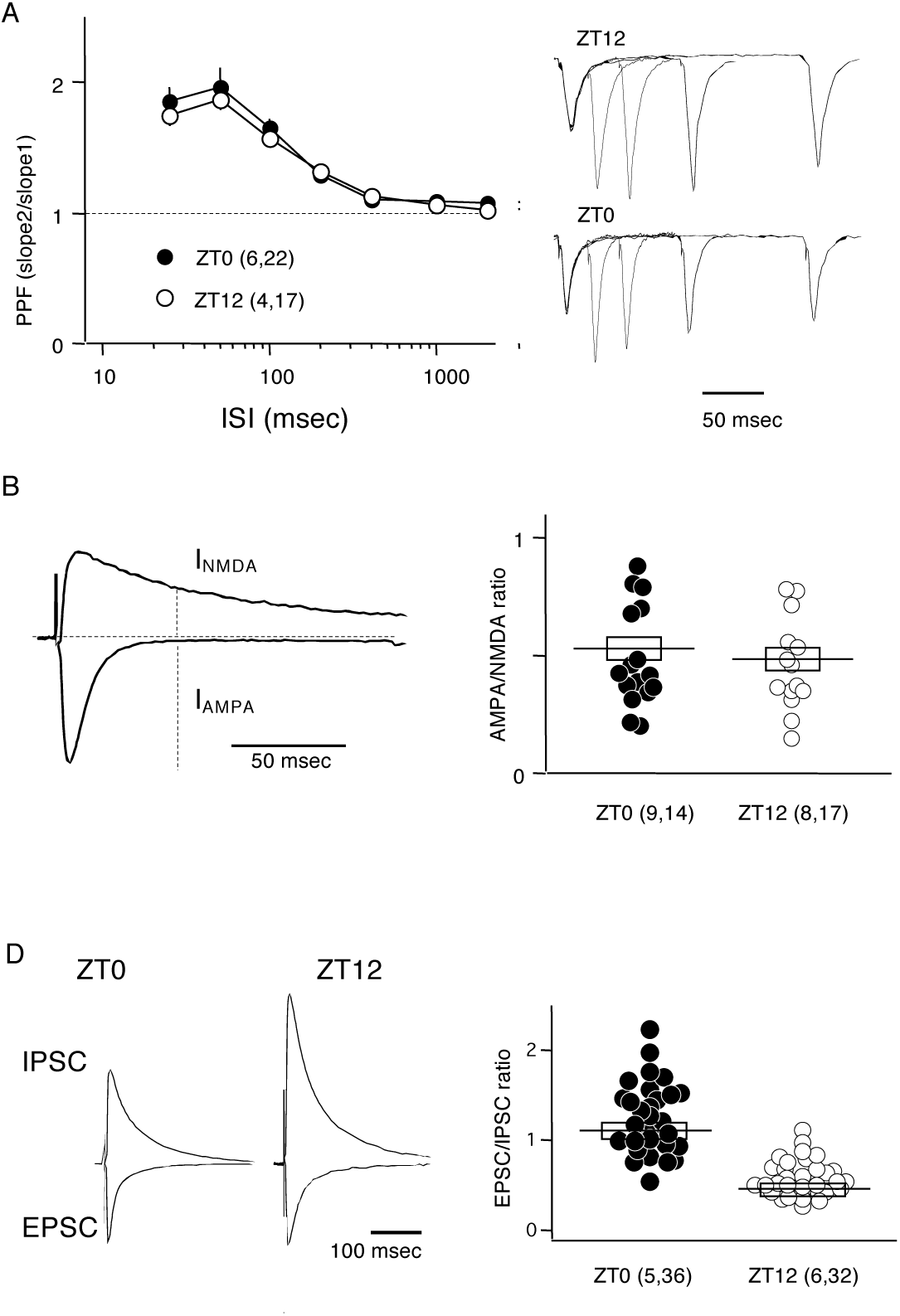
Among key LTP determinants, only daily changes in the E/I balance mirror the changes in TBS-LTP. A) Comparable paired-pulse ratio of FP slope over a range of inter-stimulus intervals (ISI: 25-2000 ms) in slices harvested at ZT0 (black circles) and at ZT12 (open circles). Superimposed traces are examples of responses evoked by paired stimulation of short intervals (25ms, 50ms, 100ms, 200ms) in a ZT0 and a ZT12 slice. B) Similar AMPA/NMDA ratio in pyramidal cells from slices harvested at ZT0 and ZT12. Traces on the left illustrate the NMDA component of the synaptic current, measured 50 msec after stimulation and recorded at Vh = +40mV (red trace), and the AMPA component, measured as the peak recorded at Vh = −70mV (black trace). The AMPA/NMDA ratio for cells from slices harvested at ZT0 (black circles) and ZT12 (open circles). C) The EPSC/IPSC ratio is in cells from slices harvested at ZT0 than at ZT12. Left: traces are example IPSCs and EPSCs (average of 4 each) recorded in two cells harvested at ZT0 and ZT12 and normalized to the EPSC peak. Right: EPSC/IPSC ratio for all cells harvested at ZT0 (black circles) and ZT12 (open circles). Horizontal lines and boxes in B and D, and data point circles in A and C, indicate average ± s.e.m. In A-C the number of mice and slices, or cells, is indicated in parenthesis.

Next, we evaluated daily changes in synaptic inhibition. Indeed, previously we have suggested that the daily changes in the ratio of synaptic excitation to synaptic inhibition (E/I ratio) that we reported in CA1 and V1 could influence the induction of plasticity (Bridi et al., 2020). In that study the frequency of miniature IPSCs -a bulk measure of synaptic inhibition in CA1-was found higher during the light cycle (Bridi et al., 2020), a result later confirmed with spontaneous IPSCs (Goode et al., 2022). Since elevated inhibition could potentially reduce LTP, it was therefore important to determine whether these daily changes in miniature and spontaneous IPSC frequency translate into changes in inhibition recruited by stimulating CA3 inputs. To that end, we computed the ratio of the excitatory and inhibitory component (E/I ratio) of the evoked compound responses. EPSCs and IPSCs were isolated by recording at different holding potentials (see methods). The results indicate that the E/I ratio of CA3-->CA1 input was significantly smaller at ZT12, consistent with an enhanced inhibition at time of the day (Figure 2C. t-test: p<0.0001).

In sum, of all three biophysical determinants examined above, only the E/I ratio changes daily and in a manner coincidental with TBS-LTP, which is consistent with the idea that an increase of the relative recruitment of inhibition limits TBS-LTP at ZT12. As a further test of this idea we examined the daily regulation of LTP induced with another type of conditioning: pairing low frequency stimulation with strong depolarization (see methods). This type of conditioning, done under voltage clamp, has the advantage that the plasticity outcome is independent of the recruitment of synaptic inhibition and cellular excitability. As shown in Figure 3A, there was no significant difference in paired-induced LTP in cells from slices harvested at ZT0 or ZT12 (MW-test: p=0.8467). This argues against a role for changes in the intrinsic capacity to support LTP mediating the daily oscillation of TBS-LTP, but it is concordant with a role of synaptic inhibition in the process. Finally, we tested whether directly blocking synaptic inhibition with bath application of the GABAa antagonist Gabazine (20μM) reduce the differences in TBS-LTP evoked at ZT0 and at ZT12. As shown in Figure 3B, in the presence Gabazine TBS-LTP was comparable at both ZT times (t-test: p=0.8028).

**Figure 3.**
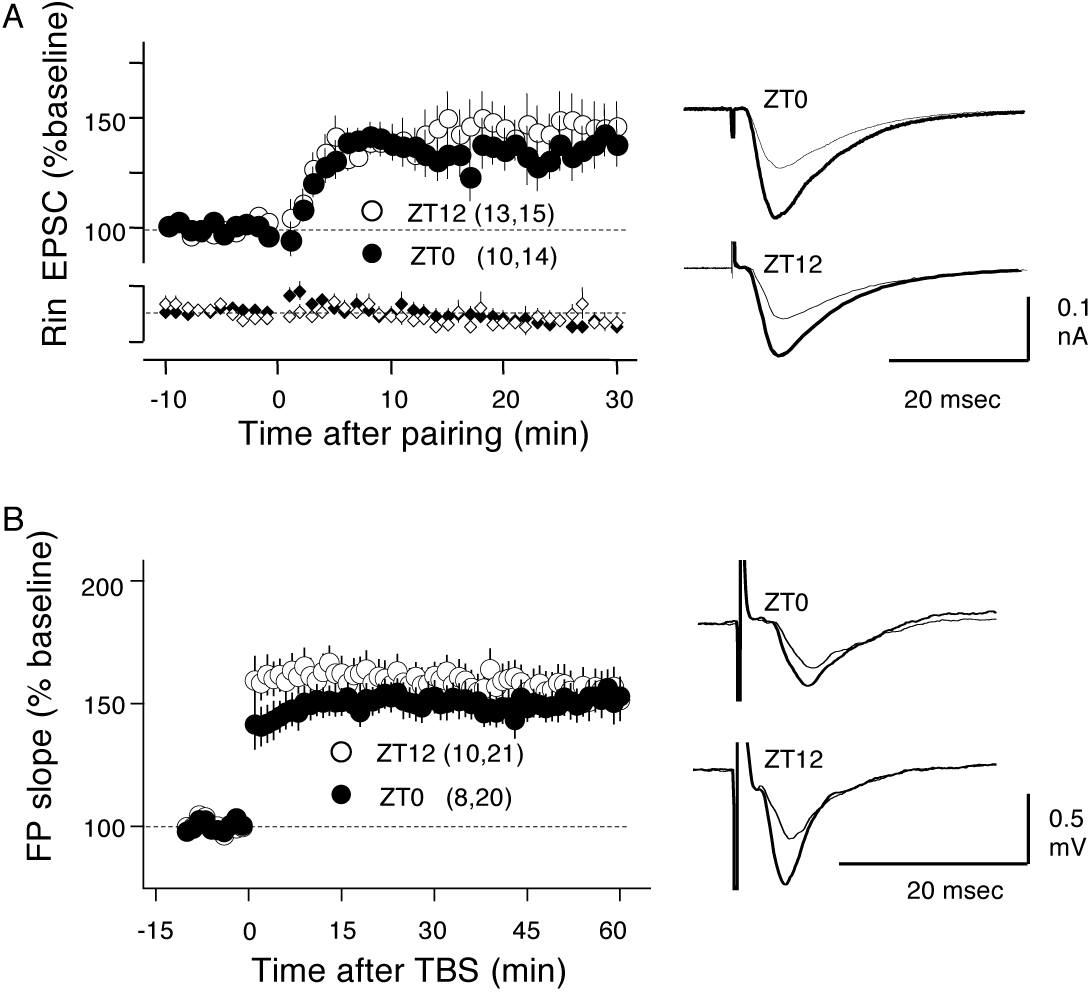
Absence of ZT-dependent differences in LTP induced in the presence of a GABAa blocker or with a pairing paradigm. A) LTP induced by a pairing paradigm (arrow: brief LFS at Vh = 0 mV see methods) in cells from slices harvested at ZT0 (black circles) and ZT12 (open circles). Left: time course of the changes in the EPSC (recorded at −70mV: top) and Rm (bottom). the number of mice and slices, or cells, is indicated in parenthesis. B) TBS-LTP induced in slices harvested at ZT0 (black circles) and ZT12 (open circles). Left: time course of fEPSP changes. Right: example traces (average of 10 consecutive responses) collected before (thin lines) and ∼50 min after TBS (thick) lines.

### Blunted TBS-LTP daily oscillations in the PS1/APP mouse model of Alzheimer’s disease

TBS-LTP in CA3→CA1 is widely used to evaluate the status of synaptic plasticity in multiple mouse models of diseases and neural conditions. It was of interest to evaluate whether changes in the daily oscillations contribute to the altered LTP magnitude documented in some of these studies. The PS1/APP mouse line was interesting because multiple studies report disparate results on LTP-TBS (reviewed by (Marchetti and Marie, 2011)). We reasoned that altered LTP oscillations might result in differences in LTP magnitude between wild types and transgenic individuals at some ZT times but not in others. To test the idea, we examined TBS-LTP in slices of adult (8 month-old) transgenic (Tg) and wild type (WT) littermates harvested at ZT0 and ZT12.

At this age cognitive deficits and alterations in NMDAR-dependent plasticity are well developed in this line (Megill et al., 2015). The results, shown in Figure 4, indicate clear differences in TBS-LTP magnitude between ZT0 and ZT12 in slices from WT but not from Tg mice. A 2-way ANOVA (Interaction: F(1,107)=3.952, p=0.0494) followed by Sidak’s multiple comparison test set at α=0.05 confirmed the significance of the results. Associated with this discrepancy in daily oscillations, the magnitude of TBS-LTP is clearly larger in WT than Tg only in slices collected at ZT12, not at ZT0 (confirmed by Sidak’s test). Thus, the TBS-LTP differences between WT and Tg markedly vary during the course of day.

**Figure 4.**
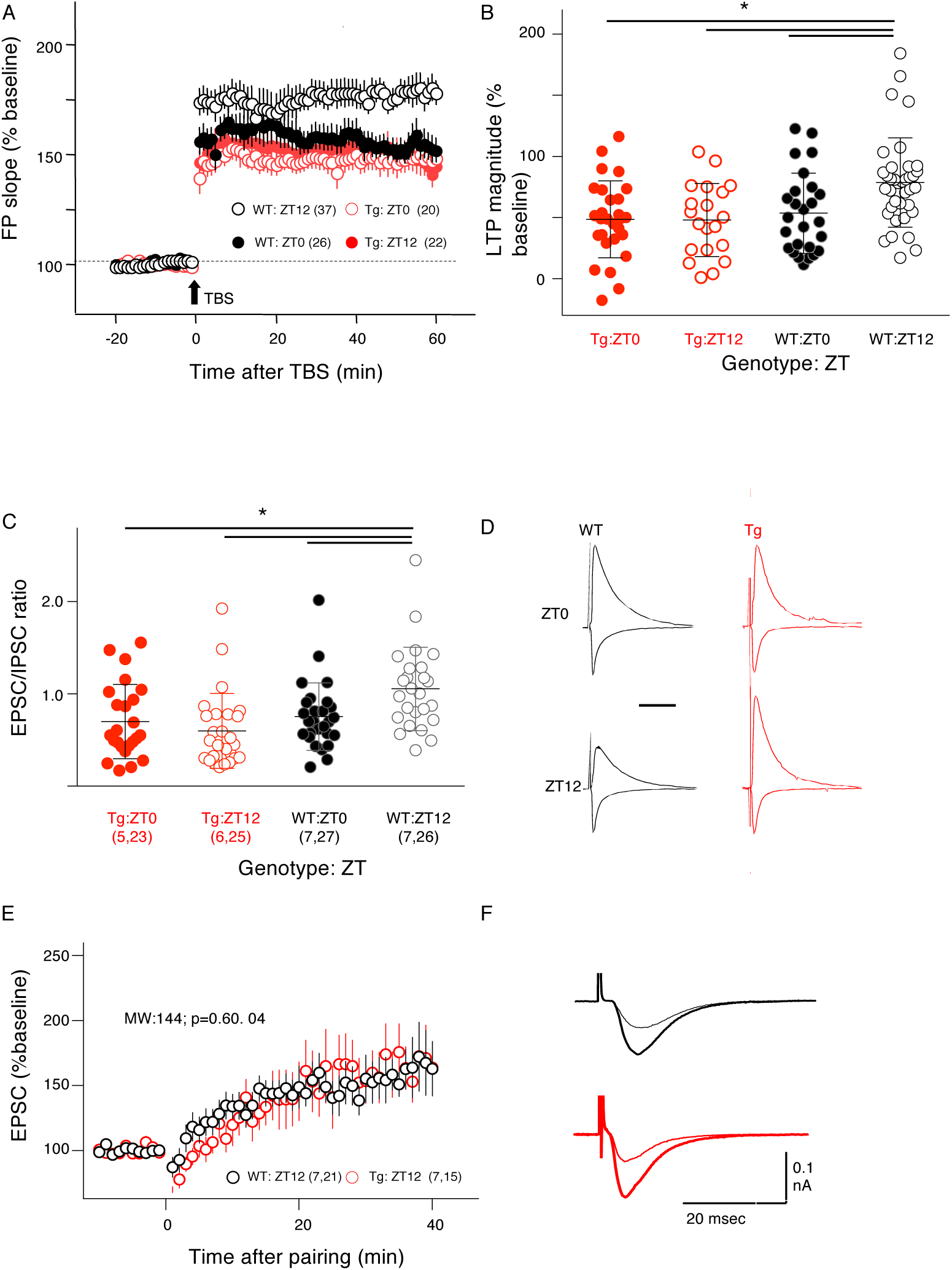
Dysregulated daily oscillations of TBS-LTP and E/I ratio in 8-month old APP/PS1 mice. A) Time course of TBS-induced changes in FP slope in slices harvested at ZT0 (filled symbols) and ZT12 (open symbols) from 8-month old transgenic APP/PS1 (Tg: red symbols) and Wild-type mice (WT: black symbols). The number of mice and slices is indicated in parenthesis. B) LTP magnitude measured 60 min after TBS in the 4 condition tested (2 genotypes, 2 ZT times). C) E/I ratio values in cells from Tg and WT mice collected at ZT0 and ZT12 (symbols as in B). D) Traces are example IPSCs and EPSCs (average of 4 each) recorded in WT and Tg cells harvested at ZT0 and ZT12 and normalized to the EPSC peak. Horizontal lines in B and C indicate average ± s.e.m, asterisks indicate Sidak’s test significance after 2-way ANOVA. Indicated in parenthesis is the number of mice and cells.

Next, we asked whether alterations in the E/I ratio daily dynamics contribute to the absence of TBS-LTP oscillations in the Tg mouse. As shown in Figures 4D and 4C, like TBS-LTP, the E/I ratio was significantly higher at ZT0 compared to in WT mice, but no such difference was observed in Tg mice (Interaction: F(1,107) = 3.952, p = 0.0494. This double correspondence between TBS-LTP and E/I ratio across ZT-time and genotype prompted us to evaluate pairing-induced LTP, as its magnitude is independent of the recruitment of inhibition. To this end, we compared pairing-induced LTP in WT and Tg cells harvested at ZT12, a time point when TBS-LTP differs significantly between genotypes. The results indicate that, unlike the case of TBS-LTP, at ZT12 the magnitude of paired-induced LTP was not different between WT and Tg (Figure 4E: MW-test: p=0.8467), suggesting a comparable capacity for LTP. Overall, these results strongly suggest that altered E/I daily dynamic is a key driver of the altered TBS-LTP observed in 8-month-old APP/PS1 mice.

The absence of daily oscillations of TBS-LTP and E/I ratio in Tg mice prompted us to question whether it relates to alterations in circadian rhythms. To that end we undertook a comprehensive evaluation of the circadian regulation of motor activity in 8-month Tg and WT mice (Figure 4. See methods). The analysis of the actograms indicates that both genotypes were equally responsive to a phase advance in the light/dark regime (Figure 5A, B. Two-way ANOVA. Interaction: F[13,112]=0.2708, p=0.9945; Days: F[1,112]=63,16; p<0.001; Genotype: F[1,112]=0.00058, p=0.9391.

**Figure 5.**
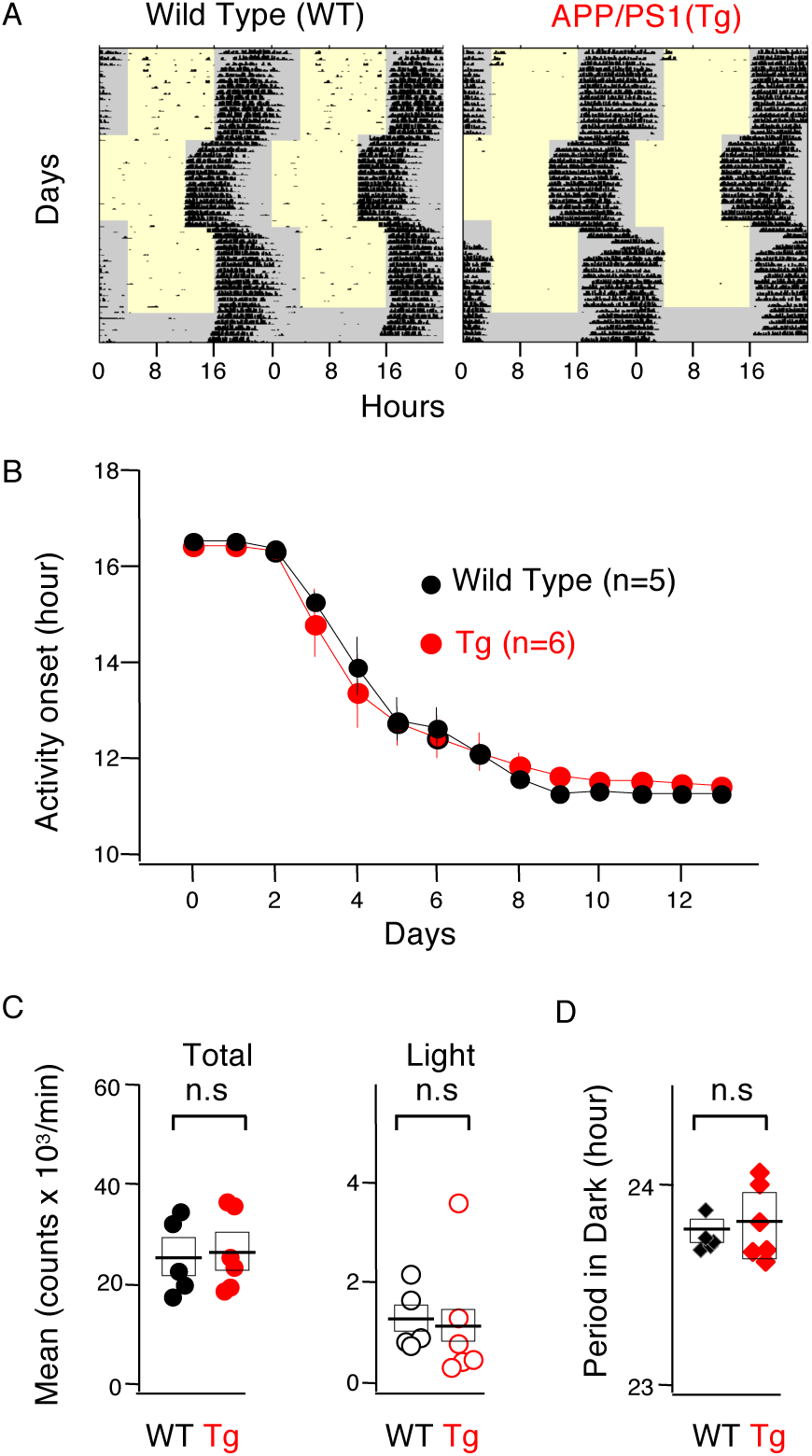
Circadian regulation of motor activity is normal in 8-month old APP/PS1 mice. Actograms were continuously recorded. By day 18 the dark cycle was advanced by 4 hours, and by day 48 the mice were kept in continuous dark. A) Example actograms in wild type (left) and transgenic APP/PS1 individuals (right). Activity is indicated as black dots. B) Phase shift response. Plotted is the time of the onset of activity in the dark vs. the day after 4-hour advance shift. Data points indicate average ± s.e.m. C) Activity in the initial period (light/dark 12:12). Left: daily total activity (light and dark); right: activity in the restive phase (light). D) Period duration (in hours) while in constant dark.

Similarly, there were no significant differences in daily activity (Figure 5C), either total activity per day (MW:p=0.329) or in the percentage of activity during the light phase (MW: p=0.0823). Finally, the period duration, measured under constant dark, was similar in both genotypes (MW-test: p=0.9719). Thus, the lack of TBS-LTP in PS1/APP mice cannot be directly attributed to altered circadian rhythms.

### Age-dependent reversal of the daily oscillation in TBS-LTP and E/I ratio

A glance comparison of the TBS-LTP and E/I ratio data presented thus far suggested a striking difference in the daily oscillations between young (4-5weeks), and adult (∼45 weeks) mice. In both cases TBS-LTP and E/I ratio change during the day, but while in young mice TBS-LTP and E/I ratio are larger at ZT0 (Figure1,2), in adults both are larger at ZT12 (Figure 4), suggesting an age dependent reversal of the daily oscillations. We hypothesized that if a developmental flip occurs, then the ZT0-ZT12 differences should be minimal at an intermediate age. To evaluate that possibility, we collected additional data from 10 weeks mice. The results indicate no significant differences between the two ZT time points for either TBS-LTP (Figure 6A; t-test: p=0.5611) or the E/I ratio (Figure 6B; MW-test: p=0.158). To visualize the developmental changes in the daily oscillations of TBS-LTP and E/I ratio, we aggregated the results of Figures 1-4 and 6 A, B and presented them in Figure 6C. This revealed a clear flip by the ∼10^th^ week of age for both, TBS-LTP and E/I ratio. In both cases, a two-way ANOVA test confirmed the interaction between age and ZT-time (Interaction for TBS-LTP: F (2, 119) = 11.93; p<0.0001. Interaction for E/I ratio: F (2, 171) = 19.17; p<0001). In the case of TBS, the analysis also revealed a significance increase with age (F (2, 119) = 10.20; p<0.0001) manifested as a sustained age-increase in LTP at ZT0 while LTP at ZT12 remained largely that the same level across the ages tested. In sum, the results suggest that changes in E/I ratio are a main determinant of changes in TBS-LTP across the day and across age.

**Figure 6.**
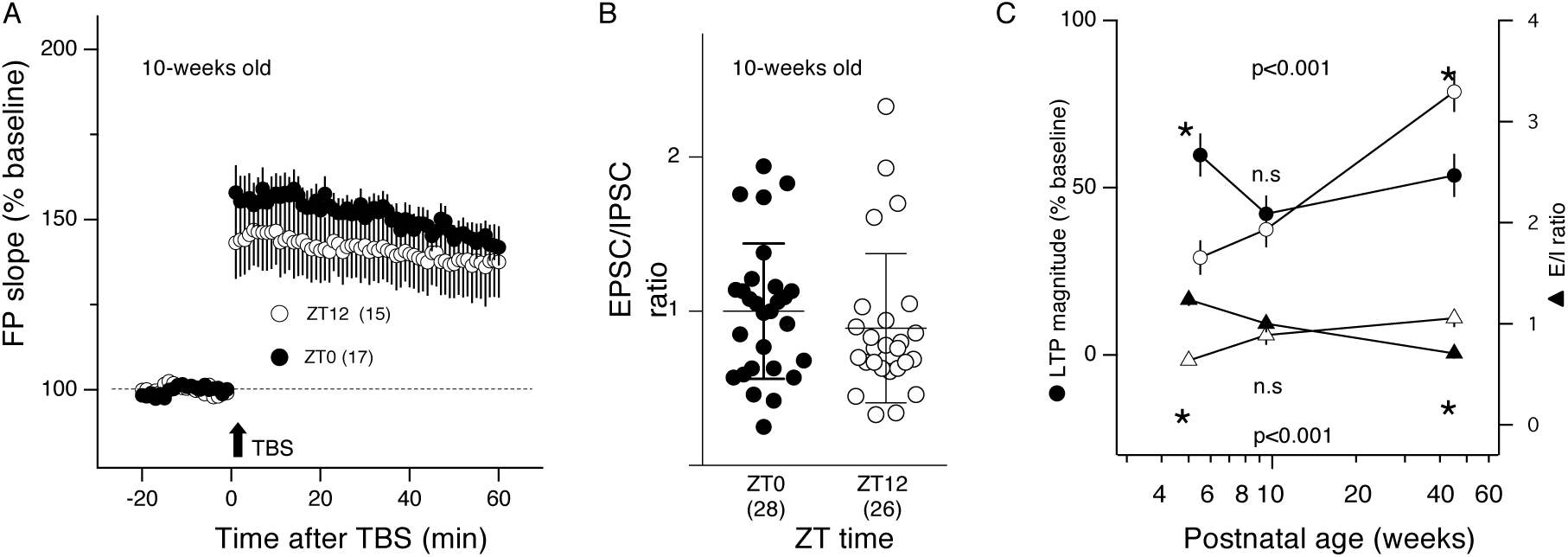
Reversal of the daily oscillations of LTP and E/I balance between 6 and 40 weeks of age. A, B) At the intermediate age of 10 weeks, the differences between ZT0 (filled circles) and ZT12 (open circles) are minimal for both TBS-LTP (A) and the E/I ratio (B). C) Course of age-dependent changes in TBS-LTP (circles, left axis) and E/I balance (triangles; right axis) in slices collected at ZT0 (filled circles) and ZT12 (open circles). The 6-week data is from figures 1 and 2; the 40-week from figure 4. In each case it is indicated the p value of the 2-way ANOVA interaction of age x ZT-time; asterisks denotes significance between ZT0 and ZT12.

## DISCUSSION

Our analysis of circadian regulation of LTP in CA1 revealed that the diminished magnitude of TBS-induced LTP across the daily light cycle primarily arises from changes in the synaptic excitation/inhibition balance. Consistent with this, TBS-induced LTP, but not LFS-induced LTD, fluctuates in parallel with the E/I ratio during the light cycle. Moreover, removing synaptic inhibition as a key determinant of LTP induction eliminates the time-of-day differences in TBS-LTP. Additional support came from the observation that in the APP/PS1 model of AD neither the E/I ratio nor TBS-LTP change during the light cycle. Notably, we also found that the daily co-changes in both the E/I ratio and TBS-LTP occur in opposite directions in slices from mature versus juvenile animals.

Although circadian regulation of LTP and its impact on behavior has been recognized for decades (for reviews see(Abel et al., 2013; Cooper et al., 2018; Lodovichi and Ratto, 2023; Snider et al., 2018), the underlying mechanisms remain not fully resolved. Our findings support a model in which the capacity to support LTP is largely constant during the day but conditions regulating its induction by high frequency stimulation vary. We propose that the excitation/inhibition (E/I) ratio is a key circadian regulator of LTP induction. In addition to the multiple correlations observed between the variations of E/I ratio and TBS-LTP across the day (also reported in (Goode et al., 2022)), we found no time-of-day differences in pairing-induced LTP -which is independent on inhibitory strength-or when TBS-LTP was induced under pharmacological blockade of synaptic inhibition. Indeed, the daily changes of E/I ratio in the face of seemingly constant NMDAR function, could readily account for the observed changes in frequency-dependent Hebbian plasticity. At ZT0, when synaptic inhibition is relatively weak, high-frequency stimulation is more likely to recruit sufficient voltage-dependent NMDAR conductance to induce LTP. In contrast, because LTD requires a lower threshold for NMDAR activation, its induction by low-frequency stimulation is expected to be less affected by daily fluctuations in the E/I ratio. In addition, the stability of LFS-LTD throughout the day is consistent with the notion that it might rely on non-ionic NMDAR functions (Dore et al., 2016; Dore et al., 2017; Nabavi et al., 2013; Stein et al., 2015). Intriguingly, it has been reported that in anesthetized rats in vivo, LFS fails to induce LTD during the dark cycle, in contrast to the present results obtained in mouse slices. The mechanisms underlying this discrepancy remain unclear.

The daily regulation of LTP by changes in GABA conductances described here for CA1 may complement a circadian modulation of LTP reported in neocortical cells, which arises from shifts in the reversal potential of GABA responses (Alfonsa et al., 2025). In addition to these GABAergic mechanisms, circadian fluctuations in fundamental membrane properties—such as intrinsic excitability (Gonzalez et al., 2023; Naseri Kouzehgarani et al., 2020)-are also likely to shape the recruitment of NMDAR conductances and, in turn, the induction of Hebbian plasticity in frequency-dependent paradigms. In addition, several molecules and processes known to influence LTP induction also undergo circadian-like regulation (for reviews, see (Cooper et al., 2018; Snider et al., 2018)). In particular, three examples—GSK3 kinase activity, synaptic BMAL1 phosphorylation, and astrocyte-derived corticosterone (Barone et al., 2023; Besing et al., 2017)—demonstrate that manipulations altering their daily dynamics also shift the circadian regulation of LTP induction. However, because TBS was the only induction protocol tested in these studies, it remains unclear whether these manipulations directly affected the synaptic capacity to support LTP or instead influenced contingencies that modulate induction, like membrane excitability, capacitance or the E/I balance. Although the daily regulation of these contingencies remains unclear, key determinants include fluctuations in cellular redox state influencing intrinsic excitability (Naseri Kouzehgarani et al., 2020) while sleep and endocannabinoid signaling modulate the E/I balance (Bridi et al., 2020).

TBS-LTP in CA1 is a widely used model for assessing altered synaptic plasticity in rodent models of various neurological conditions, including Alzheimer’s disease, schizophrenia, and autism. The time of the day regulation of LTP is a potential confound that can complicate the interpretation of differences in TBS-LTP between control and experimental animals, since experiments performed on slices harvested at different times of day are likely to yield divergent outcomes. For instance, in AD models, TBS-LTP fails to show normal daily oscillations (Carvalho da Silva et al., 2022; He et al., 2020; Li et al., 2023). Consistent with this, our data show that in APP/PS1 mouse (figure 4), TBS-LTP differs from wild type only at ZT12, but not at ZT0. Importantly, as shown here, the deficit at ZT12 appears to arise from the lack of daily fluctuations in E/I balance rather than from an intrinsic impairment in LTP, since pairing-induced LTP at ZT12 is intact in these mice. This loss of E/I-driven daily regulation of TBS-LTP may contribute to the discrepancies in the literature regarding the status of LTP in the APP/PS1 model (Li et al., 2023). An additional source of discrepancies in the literature is a developmental form of metaplasticity, whereby TBS-LTP—but not LTD—increases with age, a process that is absent in APP/PS1 mice (Megill et al., 2015). Consequently, genotype differences are expected to become more pronounced with age. Notably, and in line with this, our data revealed a selective developmental increase in both the E/I ratio and TBS-LTP, but only in slices harvested at ZT12 (figure 6). A striking consequence of these changes was a post-pubertal reversal in the time-of-day differences of the E/I ratio and TBS-LTP, which may also help explain previously reported discrepancies in the daily regulation of the TBS-LTP (Snider et al., 2018).

Specifically, TBS-LTP is larger at ZT0 than at ZT12 in juveniles, approximately equal at the end of puberty, and larger at ZT12 in adults.

In sum, our results suggest that after the completion of the initial early maturation during the first few postnatal weeks, the intrinsic cellular machinery for LTP in CA1 remains largely stable well into adulthood, with its regulation during the day depending primarily on external factors such as the E/I balance. From a methodological perspective, these findings raise a cautionary note against relying solely on TBS-LTP to assess the intrinsic capacity for Hebbian plasticity. At the same time, although the underlying mechanisms responsible for the late developmental switch of the daily regulation of E/I and TBS-LTP remain unclear, this shift may provide novel insights into how learning and memory evolve with age.

The authors declare no competing financial interests.

## Acknowledgments

Supported by NIA grants grant PO1-AG09973

